# Mitochondrial Genome Diversity across the Subphylum Saccharomycotina

**DOI:** 10.1101/2023.07.28.551029

**Authors:** John F. Wolters, Abigail L. LaBella, Dana A. Opulente, Antonis Rokas, Chris Todd Hittinger

## Abstract

Eukaryotic life depends on the functional elements encoded by both the nuclear genome and organellar genomes, such as those contained within the mitochondria. The content, size, and structure of the mitochondrial genome varies across organisms with potentially large implications for phenotypic variance and resulting evolutionary trajectories. Among yeasts in the subphylum Saccharomycotina, extensive differences have been observed in various species relative to the model yeast *Saccharomyces cerevisiae*, but mitochondrial genome sampling across many groups has been scarce, even as hundreds of nuclear genomes have become available. By extracting mitochondrial assemblies from existing short-read genome sequence datasets, we have greatly expanded both the number of available genomes and the coverage across sparsely sampled clades. Comparison of 353 yeast mitochondrial genomes revealed that, while size and GC content were fairly consistent across species, those in the genera *Metschnikowia* and *Saccharomyces* trended larger, while several species in the order Saccharomycetales, which includes *S. cerevisiae*, exhibited lower GC content. Extreme examples for both size and GC content were scattered throughout the subphylum. All mitochondrial genomes shared a core set of protein-coding genes for Complexes III, IV, and V, but they varied in the presence or absence of mitochondrially-encoded canonical Complex I genes. We traced the loss of Complex I genes to a major event in the ancestor of the orders Saccharomycetales and Saccharomycodales, but we also observed several independent losses in the orders Phaffomycetales, Pichiales, and Dipodascales. In contrast to prior hypotheses based on smaller-scale datasets, comparison of evolutionary rates in protein-coding genes showed no bias towards elevated rates among aerobically fermenting (Crabtree/Warburg-positive) yeasts. Mitochondrial introns were widely distributed, but they were highly enriched in some groups. The majority of mitochondrial introns were poorly conserved within groups, but several were shared within groups, between groups, and even across taxonomic orders, which is consistent with horizontal gene transfer, likely involving homing endonucleases acting as selfish elements. As the number of available fungal nuclear genomes continues to expand, the methods described here to retrieve mitochondrial genome sequences from these datasets will prove invaluable to ensuring that studies of fungal mitochondrial genomes keep pace with their nuclear counterparts.

## 1 Introduction

Eukaryotic evolution is a history of multiple genomes coming together. The acquisition of the mitochondria via endosymbiosis enabled new metabolic capacities, but it required the coevolution of two distinct genomes over time and created a novel dynamic (Muñoz-Gómez et al., 2015; Zachar and Szathmáry, 2017). In the vast majority of eukaryotic organisms, the mitochondrial genome (mtDNA) has been vastly reduced to encode a small number of respiratory proteins and their corresponding translational machinery (Johnston and Williams, 2016). All other ancestral mitochondrial genes were either lost or transferred to the nuclear genome, which encodes nearly all genes required for the various mitochondrial functions (Adams and Palmer, 2003). Among extant mtDNAs, there is considerable variation in specific gene content, genome structure, and idiosyncrasies of gene expression (Santamaria et al., 2007; Gualberto et al., 2014; Hao, 2022; Dowling and Wolff, 2023). Dense sampling of eukaryotic taxa is required to understand how this variation arises and its impacts on the evolution and function of both genomes.

Budding yeasts of the subphylum Saccharomycotina (hereafter, yeasts) provide a valuable model for exploring this variation further. The early sequencing of the mtDNA of the model yeast *Saccharomyces cerevisiae* provided a contrast to the picture of mitochondrial evolution that was emerging from animal studies. Whereas most animal mtDNAs were found to be highly gene-dense, small at typically under 20kb (Santamaria et al., 2007), and lacking introns, the *S. cerevisiae* mtDNA was several times larger (∼75-85kb), contained fewer genes due to lacking any of the canonical mitochondrially-encoded components of Complex I of the electron transport chain, and contained introns in several genes (Foury et al., 1998). Further studies of other eukaryotic groups confirmed that marked differences from the smaller genome seen in animals are the norm (Gualberto and Newton, 2017; Sandor et al., 2018). The addition of mtDNAs from other yeasts showed that differences in genome size were widespread and that many yeast mtDNAs still encoded a canonical Complex I (Freel et al., 2015; Xiao et al., 2017). However, the current sampling of yeast mtDNAs (Christinaki et al., 2022) remains heavily tilted towards yeasts in the order Saccharomycetales, which contains *S. cerevisiae,* and the order Serinales, which contains the opportunistic pathogen *Candida albicans* (Butler et al., 2009), but these are only two of the 12 orders in the 400-million-year-old subphylum Saccharomycotina (Shen et al., 2018; Groenewald et al., 2023) .

Yeasts have become an important model for studying the dynamics of genome evolution and, in particular, its interplay with metabolism (Scannell et al., 2011; Hittinger, 2013; Hittinger et al., 2015; Opulente et al., 2018). Nuclear genome sequences for hundreds of species across all major clades within Saccharomycotina are now available (Shen et al., 2018). However, the availability of mtDNAs for this subphylum is comparatively lacking. In this work, we demonstrate that yeast mtDNAs can be recovered from publicly available short-read genome sequencing datasets, and we more than doubled the number of available mitochondrial genomes across the subphylum to 353 mtDNAs. We show that there is considerable variation in genome size, GC content, patterns of selection, and intron content. Comparisons of gene content revealed that, while there was a major loss of Complex I in the evolution of the ancestor of the orders Saccharomycetales and Saccharomycodales, there are several additional independent losses in other orders. This dataset provides new opportunities to better understand mitochondrial evolution and its relationship to nuclear genome evolution.

## 2 Results

### 2.1 Mitochondrial Genome Sequence Rescue

To expand the availability of mtDNAs across the subphylum Saccharomycotina, we used a two-pronged approach: first searching for mitochondrial sequences in existing genome assemblies, followed by constructing new genome assemblies using assemblers specialized in generating organellar genomes from short sequencing reads. By searching for matches to existing references, we identified a treasure trove of mitochondrial sequences within the existing assemblies with sizes in the expected ranges for mtDNAs and with elevated coverage relative to the rest of the assembly, which would be consistent with the high copy number expected for the mtDNA (Solieri, 2010) (Figure 1). The success rate for extracting nearly complete mtDNAs was quite high for newer assemblies, but it was lower for older assemblies due to either lack of coverage, previously applied computational filters to remove mtDNA, or potentially the use of strains lacking mtDNA to reduce sequencing costs (Supplemental Figure 1). When raw DNA sequencing reads were readily available, reassembly by targeting mitochondrial sequences proved to be even more effective. Out of 232 species assessed via both approaches, 19 were best assembled within the nuclear assembly, whereas 212 were best completed via reassembly (38 by plasmidSPAdes (Antipov et al., 2016) and 174 by NOVOPlasty (Dierckxsens et al., 2017)). After reducing the mitochondrial genome assemblies to the best representative for each species (Supplemental Table 1), the number of Saccharomycotina species with mtDNAs available increased from 132 (Christinaki et al., 2022) to 353, which included dramatically improved representation in several clades (Figure 2). Many of these mtDNAs were assembled as a circle, but a small number of assemblies remained fragmented, which resulted in missing portions with contig breakpoints that occasionally overlapped annotated genes. The pipeline for searching existing genome assemblies for mitochondrial sequences is available here: https://github.com/JFWolters/IdentifyMitoContigs.

**Figure 1.**
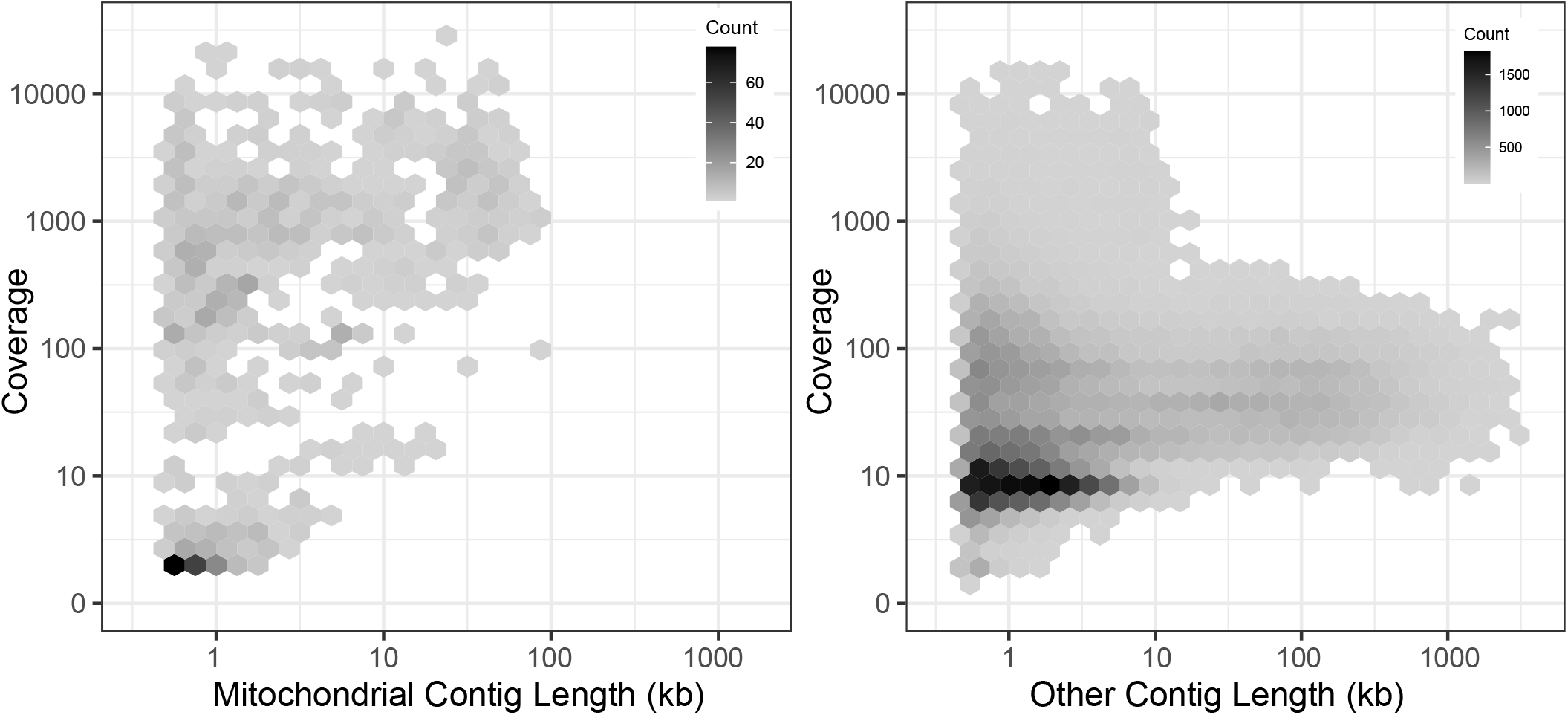
Mitochondrial Contig Profile. The coverage and length profile of contigs from 196 assemblies newly sequenced in (Shen, et al., 2018) that were flagged as putative mitochondrial contigs versus all other contigs is displayed (log_10_ scaling). The most useful mitochondrial contigs generally have a profile of elevated coverage with sizes between 10 and 100 kb, a combination rarely found in other contigs, although strict diagnostic cutoffs are not evident. Many poor-quality putative mitochondrial contigs were found in nuclear genome assemblies, but these were not present in mitochondrially-focused reassemblies.

**Figure 2.**
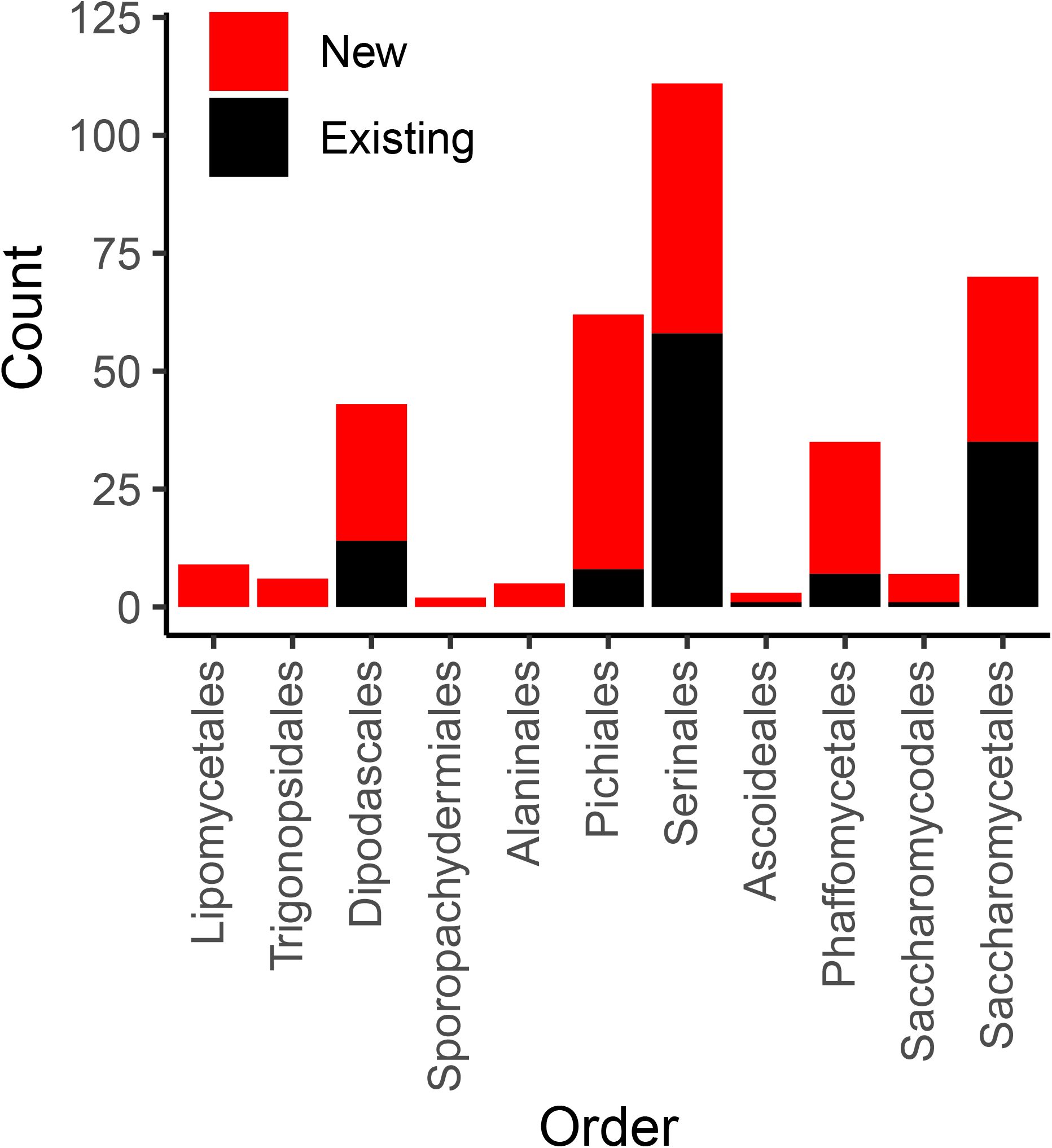
Mitochondrial Genome Counts by Taxonomic Order. The count of genomes for both newly added and existing genomes from public repositories are displayed according to taxonomic order (classifications recently described by Groenewald, et al. 2023). For nearly all orders, a majority of genomes are new (barring Saccharomycetales (35 new versus 35 existing) and Serinales (53 new versus 58 existing)).

### 2.2 Phylogeny and Genome Characteristics

We constructed a phylogeny of yeast mtDNAs based on concatenation of the core protein-coding genes (Figure 3). Overall concordance with the existing nuclear phylogeny was reasonably high (normalized Robinson-Foulds distance 0.24 between matched subtrees). Placement of the recently described (Groenewald et al., 2023) taxonomic orders (previously designated as major clades (Shen et al., 2018)) was consistent between the phylogenies, barring three exceptions: two *Trigonopsis* species grouped closer to Lipomycetales than other Trigonopsidales; the Alaninales were paraphyletic with respect to the Pichiales, rather than forming a single monophyletic outgroup; and the placement of the fast-evolving lineage of *Hanseniaspora* (order Saccharomycodales) was uncertain due to the long branch at the root of this order. A similar inconsistency was observed in prior phylogenetic analysis where *Hanseniaspora* mtDNAs clustered with the order Serinales (Christinaki et al., 2022). The uncertainty in the placement of the fast-evolving lineage of *Hanseniaspora* is likely due to long branch attraction (Bergsten, 2005). Thus, in Figure 3, we have displayed results from a tree-building run that recovered the order Saccharomycodales as monophyletic, as expected from the genome-scale nuclear phylogeny (Shen et al., 2018). Within taxonomic orders, groupings of genera were highly congruent with the genome-wide species phylogeny, but some inconsistencies remained in the placements of genera. For example, *Eremothecium* mtDNAs appeared as an outgroup to other Saccharomycetales, rather than grouping with *Kluyveromyces* and *Lachancea* as expected. Overall, we conclude that the observed mtDNA phylogeny generally tracked the species phylogeny and was not consistent with widespread introgressions or horizontal gene transfer (HGT) of protein-coding genes across long evolutionary distances.

**Figure 3.**
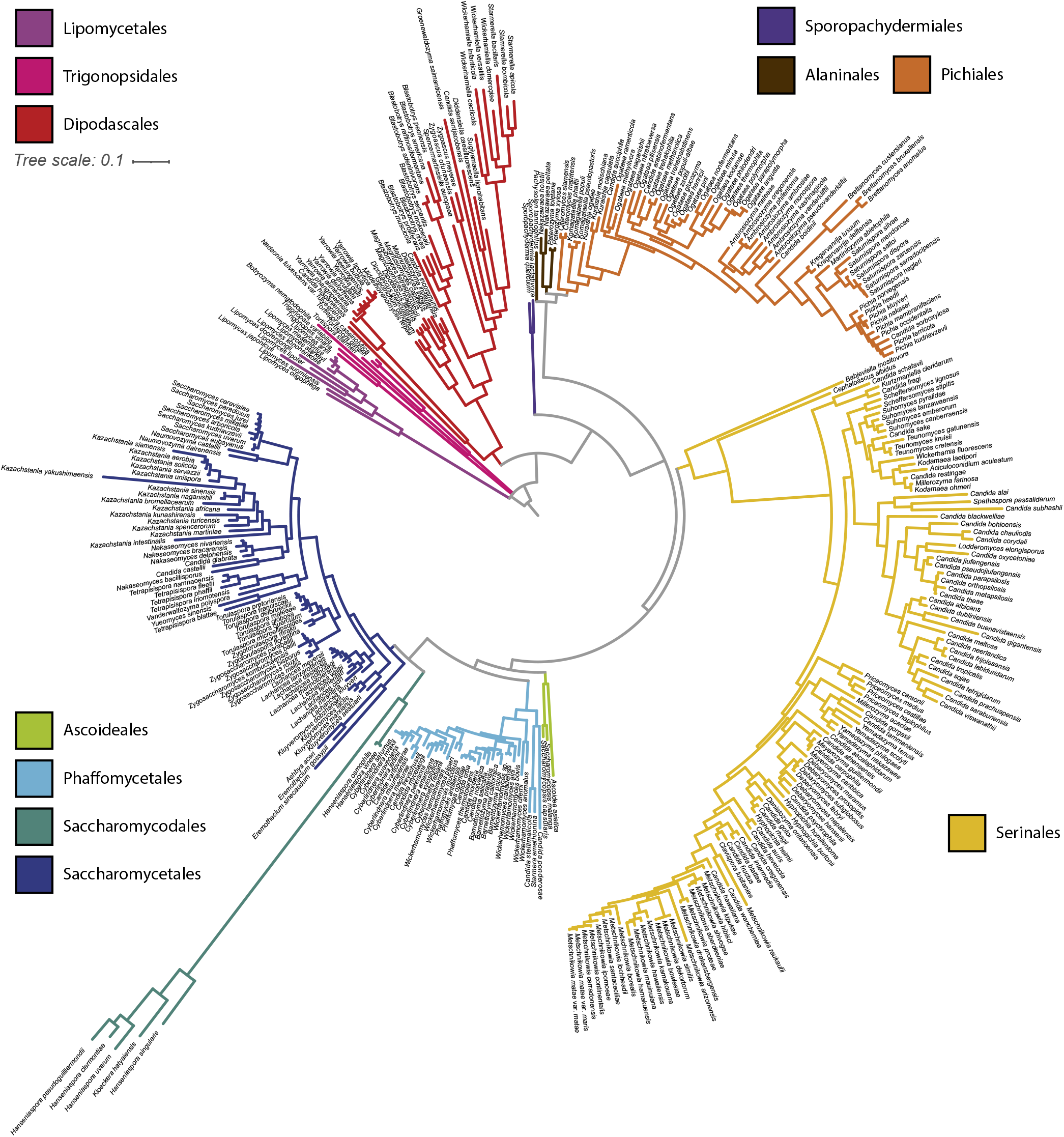
Mitochondrial Phylogeny of 353 Budding Yeasts. A phylogenetic tree was built from the protein sequences of the core protein-coding genes shared by all 353 budding yeast species analyzed (*COX1, COX2, COX3, ATP6, ATP8, ATP9*, and *COB*). Branches are colored based on taxonomic order.

Analysis of mitochondrial genome content suggested that all mtDNAs likely retain the complete set of core respiratory genes, including: the Complex IV components encoded by *COX1*, *COX2*, and *COX3*; the complex III component encoded by *COB*; and the ATP synthase components encoded by *ATP6*, *ATP8*, and *ATP9* (Figure 4A). The absence of some of these genes from a small number of assemblies was generally due to the assembly being fragmented or the annotation being manually removed due to issues with gene annotation (see Methods). In contrast, the mitochondrially-encoded components of the canonical Complex I (encoded by *NAD1*-*NAD6* and *NAD4L*) were surprisingly absent in several mtDNAs that otherwise appeared to be complete (Figure 4B). These genes are generally present in the mtDNAs of most fungi (Sandor et al., 2018) but were known to be absent in the orders Saccharomycetales and Saccharomycodales (Freel et al., 2015; Christinaki et al., 2022); indeed, our analysis is consistent with a major loss event in the common ancestor of these lineages. However, we also observed a single species lacking these genes in the order Dipodascales, *Nadsonia fulvescens* var. *fulvescens*, which is consistent with their absence in the related species *Nadsonia starkeyi-henricii* (O’Boyle et al., 2018) that was not included in this dataset, as well as a novel single-species loss event in the order Pichiales for *Ogataea philodendra*. More strikingly, there were multiple independent losses within the order Phaffomycetales, including a single loss in the ancestor of *Candida ponderosae*, *Starmera amethionina*, and *Candida stellimalicola*, as well as potentially independent losses for *Wickerhamomyces pijperi* and *Cyberlindnera petersonii*. The distribution of the ribosomal protein encoded by *RPS3* was extremely patchy (Figure 4C). *RPS3* was not universally present in any taxonomic order, but all species in the dataset from the orders Serinales, Lipomycetales, and Sporopachydermiales lacked this gene.

**Figure 4.**
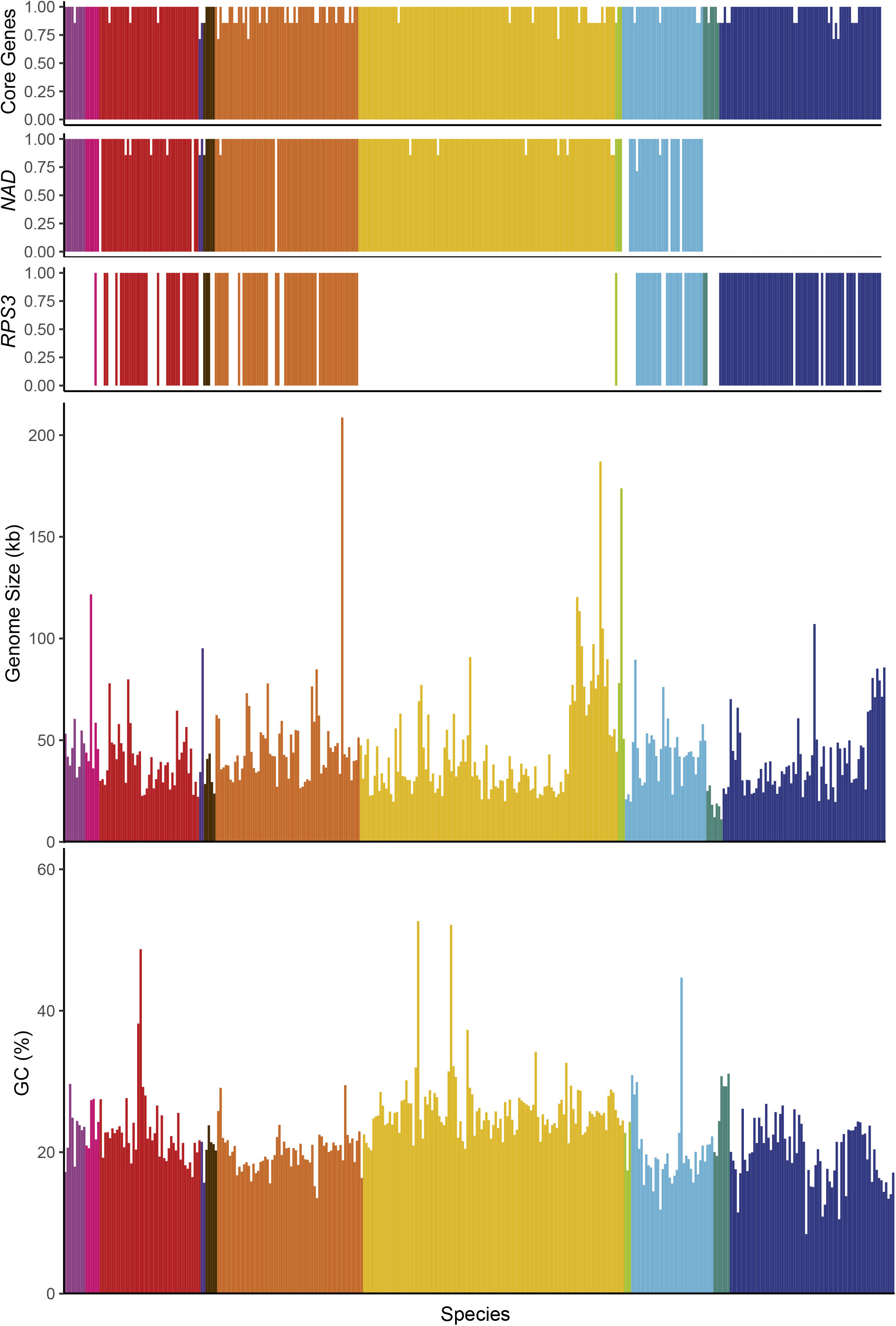
Genome Characteristics. Genome characteristics are displayed and colored according to taxonomic order and placed based on position in the phylogenetic tree (left to right from Lipomycetales to Saccharomycetales, see Figure 3). The proportion of genes found in each genome are shown for: A) core genes (*COX1, COX2, COX3, ATP6, ATP8, ATP9*, and *COB*), B) Complex I genes (*NAD1-NAD6*, and *NAD4L*), and C) the *RPS3* gene encoding a ribosomal protein. Genome sizes (D) and GC content (E) are indicated; both maintain a fairly limited range across the subphylum with a handful of extremes present across multiple taxonomic orders.

Despite similarities in gene content, genome size varied wildly at the extremes. *Pichia heedii* exceeded the previously largest observed Saccharomycotina mtDNA at 209,444 bp (versus the previous record of 187,024 bp in *Metschnikowia arizonensis* (Lee et al., 2020)), while the smallest observed mtDNA was *Hanseniaspora pseudoguilliermondii* at 11,080 bp (versus the previous record of 18.8 kb in *Hanseniaspora uvarum* (Pramateftaki et al., 2006)) (Figure 4D). The precise sizes of some mtDNAs were difficult to assess because not all assemblies were strictly complete, and short reads were not always capable of resolving genome structure reliably. The mtDNAs over 100 kb were typically more than double the size of any closely related species. Despite these observed extremes, the genome size of most species stayed within a range from approximately 20kb to 80kb (median size 39 kb, mean size 44 kb, standard deviation 23 kb). While this size variation is considerable in comparison with animal mtDNAs (Santamaria et al., 2007), it is within the ranges observed for other fungal mtDNAs (Sandor et al., 2018) and relatively low compared to plant mtDNAs (Gualberto and Newton, 2017).

The majority of species had similar GC content with a small number of outliers (Figure 4E). The average GC content was low (mean GC 22.5%, standard deviation 5.2%). Unusually high GC contents were sporadically placed around the phylogeny, including *Candida subhashii* (52.7%) and *Candida gigantensis* (52.1%) in the order Serinales, *Magnusiomyces tetraspermus* (48.7%) in the order Dipodascales, and *Wickerhamomyces hampshirensis* (44.7%) in the order Phaffomycetales. The lowest value observed was for *Tetrapisispora blattae* at 8.4% (order Saccharomycetales), which was close to lowest value of 7.6% previously observed in *Saccharomycodes ludwigii* (Nguyen et al., 2020a), which was not included in this dataset. Expansions of AT-rich intergenic regions have previously been reported to drive increases in genome size, which could drive a correlation between genome size and GC content. We found that, while this trend may be true in some groups, the overall correlation between genome size and GC content was poor and not significant after phylogenetic correction (r=-0.11, p-value 0.03; phylogenetically corrected r=-0.47, p-value 0.1, Supplemental Figure 2). Among the genomes over 100kb, the average GC content (22.8%) was close to the global average. *Nakaseomyces bacillisporus* may have driven prior correlations within smaller scale analyses of the order Saccharomycetales (Xiao et al., 2017) due its unusually large size (107 kb) and low GC content (10. 9%), but this relationship does not appear to be strong across the expanded dataset. If expansions of intergenic regions drive size variation between distant species (Hao, 2022), then they likely do so in a GC-independent manner.

### 2.3 Aerobic Fermenters Lack Evidence for Relaxed Purifying Selection

Metabolic strategies vary greatly among yeasts with regards to fermentation and respiration, which has been proposed to impact selection pressures on mitochondrial genes (Jiang et al., 2008). While many yeasts strongly respire fermentable carbon sources, such as glucose, there are many specialized yeasts, including most famously *S. cerevisiae*, that have developed metabolic strategies to preferentially ferment glucose and repress respiration, even in aerobic conditions (Merico et al., 2007; Rozpędowska et al., 2011; Hagman et al., 2013; Dashko et al., 2014; Hagman and Piškur, 2015). These aggressive fermenters are commonly said to exhibit Crabtree/Warburg Effect and are referred to as Crabtree/Warburg-positive (Diaz-Ruiz et al., 2011; Pfeiffer and Morley, 2014; Hammad et al., 2016). Given the relative disuse of respiration by this lifestyle, we hypothesized that the mitochondrially-encoded genes of Crabtree/Warburg-positive groups would exhibit elevated rates of non-synonymous substitutions due to relaxed purifying selection. Prior analysis of a limited set of species in the order Saccharomycetales had supported this model (Jiang et al., 2008).

To test the generality of this hypothesis, we determined the ratio of non-synonymous to synonymous substitution rates (ω) among groups at roughly the genus level (see Methods) across the phylogeny (Figure 5, Supplemental Table 2). We expected that ω would be highest in *Saccharomyces* and in related yeasts in the order Saccharomycetales that had undergone a whole-genome duplication (Marcet-Houben and Gabaldón, 2015; Wolfe, 2015) and were known to be strong fermenters, such as *Kazachstania* and *Nakaseomyces* (Hagman et al., 2013). Surprisingly, we observed that ω varied greatly within taxonomic orders, with many groups exceeding the values observed for *Saccharomyces*. Indeed, the highest values were found in the order Dipodascales for yeasts in the *Wickerhamiella*/*Starmerella* clade and the grouping of yeasts most closely related to that clade (referred to as “Other Dipodascales” in Figure 5). The observed values for this clade are unlikely to be an artifact caused solely by long branch-lengths because the genus with the longest branch-lengths in the phylogeny (*Hanseniaspora* in the order Saccharomycodales) exhibited relatively moderate values. Within the order Saccharomycetales, we observed a general trend towards higher ω among yeasts that underwent the whole-genome duplication. The genus *Saccharomyces* followed this trend to some extent (genus mean ω 0.09 versus global mean 0.061), but this result was primarily driven by a single gene, *ATP8,* which had the highest value observed for all genes and groups and was driven by high values on the branches leading to *S. paradoxus and S. arboricola* (0.355). When this gene was excluded, the remaining genes defied the trend (0.046). *ATP8* is highly conserved between *S. cerevisiae* strains (Wolters et al., 2015), which suggests inter- and intra-specific patterns of variation can differ greatly. Given the high ω values for many yeasts not known to be Crabtree/Warburg-positive and the relatively low ω for most *Saccharomyces* genes, we conclude that our much-expanded dataset does not support the previously proposed model of pervasive relaxed purifying selection on the mitochondrially-encoded genes of aerobic fermenters.

**Figure 5.**
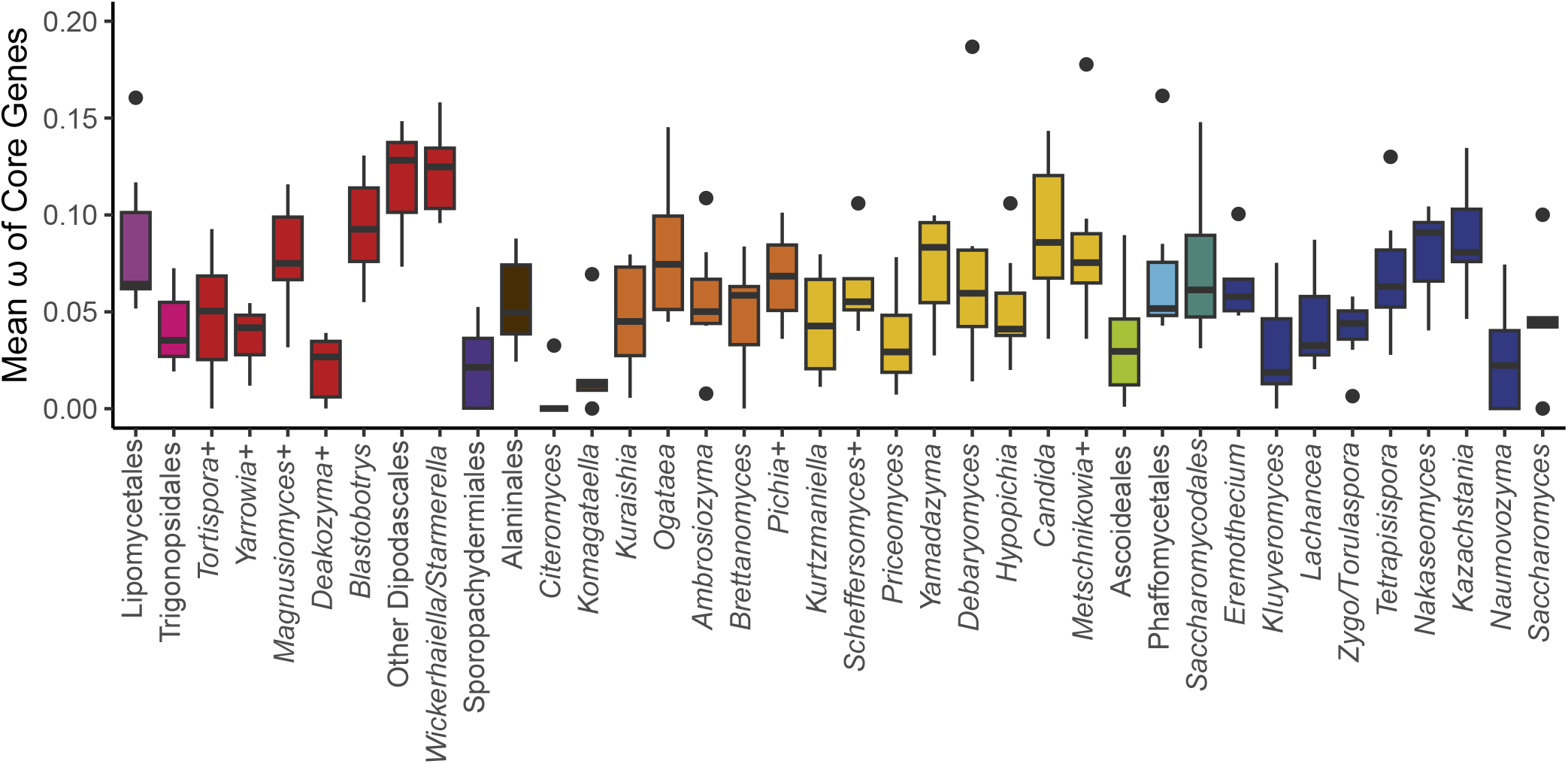
Mean ⍰ of Core Genes. The ratio of non-synonymous to synonymous substitution rates for each of the core protein-coding genes was calculated for groups across the phylogeny (+ indicates that additional closely related species that are not currently classified in that genus were included, see Table S1). The box and whisker plots show the distribution of ω among genes within each group (boxes centered at median encompassing the interquartile range, whiskers up to 1.5 times the interquartile range, and outlier genes shown as individual datapoints). Two extreme outlier genes were omitted from the graph: *ATP8* for *Saccharomyces* (0.355) and *COB* for *Kurtzmaniella* (0.250). Groups with aerobic fermenters, such as *Saccharomyces*, *Kazachstania*, and *Nakaseomyces*, do not exhibit significantly elevated ratios relative to the rest of the subphylum.

### 2.4 Evidence for Horizontal Transfer of Mitochondrial Introns Across Orders

Mitochondrial introns vary widely in yeasts, largely due to sporadic gains and losses (Xiao et al., 2017). Intron-encoded homing endonucleases are thought to drive intron turnover and potentially HGT of introns between species (Lang et al., 2007; Wu and Hao, 2014). The highest numbers of introns were observed in *Magnusiomyces* (mean 18 introns per species versus global mean 5.4 Supplemental Table 3), *Metschnikowia* (10.7), and *Yarrowia* (10.5, including other closely related anamorphic species that have yet to be reassigned to this genus). The lowest values were observed in *Eremothecium* (0.33) and *Deakozyma* (0.5), both of which included species that were completely free of introns. Nearly all introns were encoded within *COX1* (55.3%), *COB* (30.1%), or *NAD5* (7.4%); the remaining genes had <2% each. The small range of gene targets is consistent with intron homing by endonucleases transferring introns, including by HGT, to a limited range of target sites.

We identified potential intron HGTs based on BLAST comparisons of all mitochondrial introns observed using a conservative threshold to classify introns as unique, shared within a group (identical groupings as for the selection analysis above), shared within and between groups, or solely between groups (>50% of maximum possible bit score, Figure 6A). Most introns observed did not share high sequence similarity to introns from other species (65.6%), while most of the remainder were shared within a group (30%). A small number were shared across groups, and this phenomenon was especially common in the order Saccharomycetales (Figure 6B). Clustering the introns based on pairwise BLAST hits generated 271 clusters of related introns (Supplemental Table 3). Only a single cluster contained introns that were found within different genes due to homology between *Metschnikowia mauinuiana NAD2* intron 1 and *COX1* intron 1 from the same species and from *Metschnikowia hawaiiensis* (Figure 6C). *NAD2* is duplicated in *Metschnikowia mauinuiana*, but only one copy has been colonized by this intron; however, the second copy contains a 560-bp duplication identical to the 3’ end the intron. Thus, *M. mauinuiana NAD2* intron 1 may be misannotated and may instead be a 3’ terminal element that could be translated as an extension of the upstream gene; a similar phenomenon has been observed for *COX2* and other genes in *Saccharomyces* (Peris et al., 2017). *M. mauinuiana COX1* intron 1 had homology to the reverse transcriptase encoded in intron 1 of *S. cerevisiae COX1*; however, *M. mauinuiana NAD2* intron 1 appeared to be truncated, which disrupts the intronic open reading frame. Thus, *M. mauinuiana NAD2* intron 1 may better be thought of as an example of how a 3’ terminal element may be formed by an intronic mobile element acquiring a novel insertion site. The high number of introns in these species may be increasing the odds of such events in this genus, which has been speculated to have the strangest mitochondrial genomes (Lee et al., 2020).

**Figure 6.**
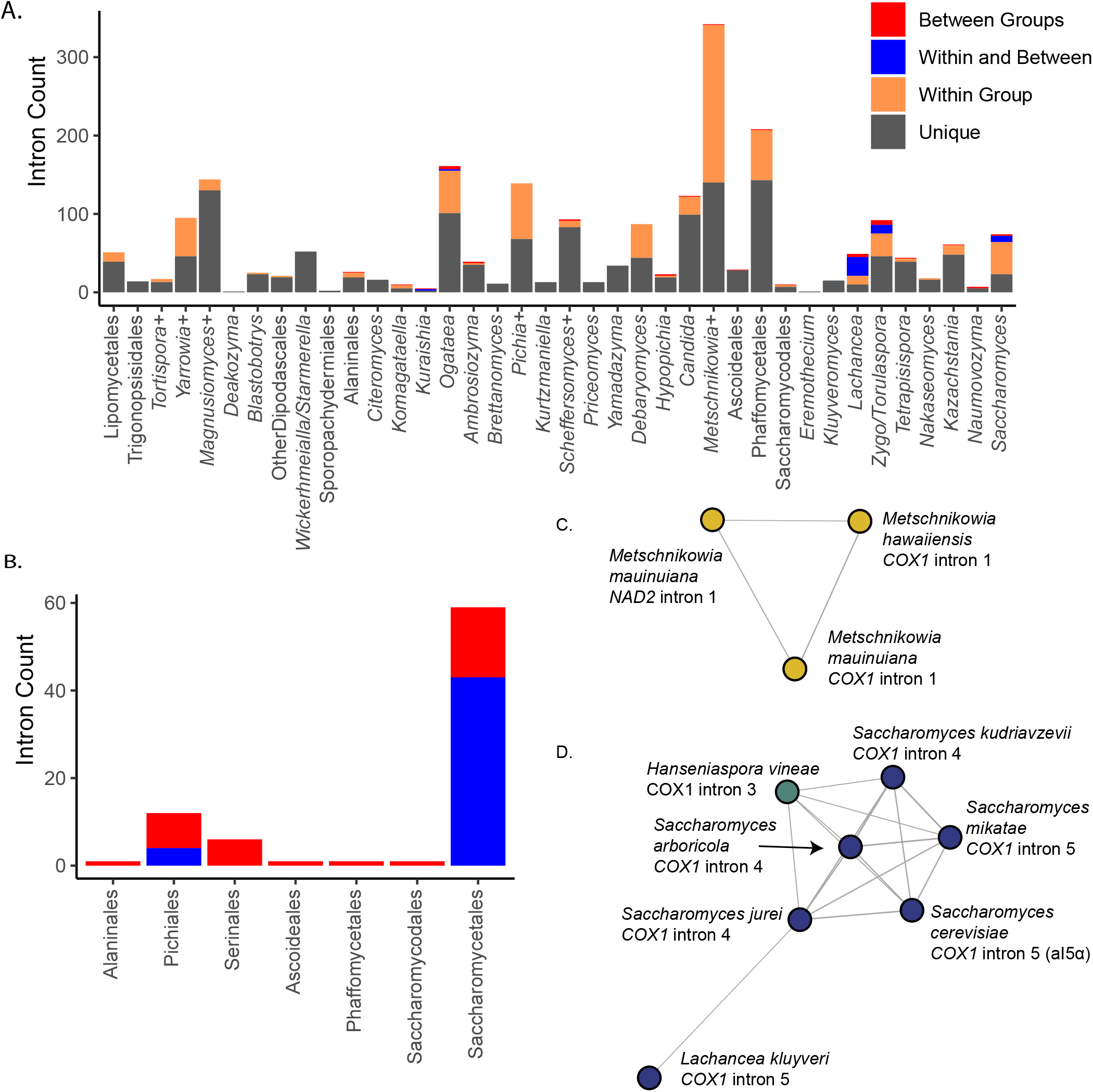
Intron Diversity. A) Introns were classified based on pairwise BLAST hits as unique to that species, present in multiple species of the group, shared within and between groups, or only between groups. The counts of introns in each category within each group are displayed. B) The counts of introns in each taxonomic order that were shared or found only between groups are displayed. Orders not listed had no introns in these categories. C) Introns were clustered based on shared BLAST hits, and the single cluster containing hits shared across multiple genes is displayed. Nodes are colored based on taxonomic order as in Figure 2 (all Serinales). D) A cluster of introns is displayed that spans the orders Saccharomycetales and Saccharomycodales, including *Saccharomyces* spp., *Lachancea kluyveri*, and *Hanseniaspora vineae*. Nodes are colored based on taxonomic order as in Figure 3.

We observed 22 clusters that contained introns spanning multiple groups, including four that contained introns spanning multiple orders (Figure 6D, Supplemental Figure 3); these clusters are the top candidates for HGT events in our dataset. For example, the fifth intron of *COX1* from *S. cerevisiae* (sometimes referred to as aI5α) shared homology with several *Saccharomyces COX1* introns, as well as *Hanseniaspora vineae COX1* intron 3 (Figure 6D). This cluster of introns may also include *Lachancea kluyveri COX1* intron 5, but this connection was only supported for *Saccharomyces jurei COX1* intron 4 (Figure 6D). All other *H. vineae COX1* introns (order Saccharomycodales) shared limited homology to introns within the order Saccharomycetales, but it was well below our cutoff; since they shared no clear homology to other *Hanseniaspora* introns, these are also candidates for HGT, albeit more tentative ones. Two of the four clusters with evidence of cross-order HGT involved introns from *Hypophichia burtonii*, which suggests that this species may contain several highly active intronic mobile elements. Interestingly, this lineage also appears to have been an HGT donor of nuclear-encoded genes for utilization of the sugar galactose (Haase et al., 2021). We conclude that homology in homing endonuclease target sites likely enables the HGT of these selfish elements, even across large phylogenetic distances, at least in rare cases.

## 3 Discussion

As high-throughput sequencing revolutionized genomics, advances in yeast mitochondrial genomics were initially delayed. Early high-throughput datasets generated only partial sequences, potentially due to biases against AT-rich sequences (Chen et al., 2013; Ross et al., 2013). Advances in methodology led to large numbers of *S. cerevisiae* mitochondrial genomes being sequenced in tandem with their nuclear genomes (Strope et al., 2015). More recently, even very large population datasets produced mitochondrial genomes concurrently with the nuclear genomes (De Chiara et al., 2020). Prior to this study, these advances had not yet come to bear for large species-rich datasets, with targeted post-hoc searches of published assemblies yielding limited numbers of additional mtDNAs (Christinaki et al., 2022). Here, we have demonstrated that, even for short-read-only datasets, it is possible to extract high-quality mitochondrial genomes with a high success rate from datasets originally collected for nuclear sequencing. As yeast genomics progresses further, the mitochondrial component need not be an afterthought.

Despite these advances, limitations remain. Mitochondrial genome structure is complex and not always readily solvable through short reads alone. For example, the *S. cerevisiae* mtDNA maps genetically as circular, but the predominant molecular form is a linear concatemer of multiple genome units (Solieri, 2010). Other species exhibit true linear forms, including *C. albicans* (Gerhold et al., 2010), or even have capping terminal inverted repeats as seen in *H. uvarum* (Pramateftaki et al., 2006). Long-read sequencing technologies are a promising avenue to obtain not only complete mtDNAs, which short reads alone failed to provide for many species, but also to resolve complex genome structures by generating reads longer than a single genome unit in length. This strategy has already been successful at investigating large-scale deletion mutations in *S. cerevisiae* (Nunn and Goyal, 2022). However, specialized assemblers, similar in principle to those used here for reassembly of the short reads, will be needed because current long-read assemblers, such as canu (Koren et al., 2017), frequently misassemble circular-mapping genomes (Wick and Holt, 2019).

The most striking variation seen among the mtDNAs is the complete loss of canonical Complex I in Saccharomycetales, Saccharomycodales, and several additional lineages across the phylogeny. In *S. cerevisiae*, the acquisition of genes encoding a multi-unit alternative NADH:ubiquinone oxidoreductase facilitated this loss (Luttik et al., 1998; Kerscher, 2000), albeit at the cost of a loss in potential proton motive force. The mechanisms that allowed for this loss in the other independent events are currently unclear, but they suggest that multiple species may also have potentiating factors that could facilitate loss. Canonical Complex I is encoded by both nuclear and mitochondrial genes, but these nuclear genes were concomitantly lost in *S. cerevisiae* with the mitochondrial genes. If the same pattern persists across all independent loss events, then it may be possible to identify unknown genes related to Complex I that were also lost in tandem.

Originally, we hypothesized that preference for aerobic fermentation would be a major factor driving mitochondrial genome variation. Previously, it had even been hypothesized to play a significant role in the loss of Complex I as *Brettanomyces* species were the only others known to be Crabtree/Warburg-positive but still encode a canonical Complex I (Freel et al., 2015). Given that multiple losses of Complex I were observed in species not known to be Crabtree/Warburg-positive and given the lack of evidence for relaxed purifying selection in aerobic fermenters, it is not evident that this shift in metabolism is a major driver of mitochondrial gene evolution. An important caveat is that the methodology employed here may be limited by current datasets on the distribution of aerobic fermentation, which extrapolate from only a handful of well-characterized species. For example, the *Wickerhamiella*/*Starmerella* clade merits further attention due to the high rates of non-synonymous variation observed and potential environmental preferences for sugar-rich environments in this group (Gonçalves et al., 2020). Additionally, estimating selection at the group level may obscure patterns of selection that vary more between closely related species than between groups, as previously observed for *Lachancea* species (Freel et al., 2014). Focusing on selection pressures at the level of individual genes may also be more illuminating. The ω rates varied more for comparisons for the same gene across groups (mean variance 0.0018) than for comparisons of different genes within groups (0.0014). For example, while *ATP9* is the most conserved gene within *Saccharomyces*, it is the least conserved in *Nakaseomyces*. If aerobic fermentation does play a role, it may relax selective pressure on some genes but increase purifying selection for others.

Mitochondrial introns may also serve an important role in shaping mitochondrial gene evolution. Homing endonucleases, which are encoded within mitochondrial introns or in downstream open reading frames at the 3’ end of mitochondrial genes, have been shown to modify sequences adjacent to the insertion site (Repar and Warnecke, 2017; Xiao et al., 2017; Wu and Hao, 2019). Transfers between groups, and potentially between orders, may introduce non-synonymous variation due to co-conversion of flanking sequences during insertion. We observed a large proportion of unique introns in our dataset, which is consistent with high rates of intron turnover underlying presence/absence variation. However, we have likely underestimated the true proportion of introns shared within groups due to the stringent criteria applied and the rapid decay of detectable sequence homology due to high mtDNA mutation rates (Sharp et al., 2018). Mitochondrial introns have been known to jump between different kingdoms between the symbiotic components of lichens (Mukhopadhyay and Hausner, 2021). Certain ecological conditions, such as coculture of *Saccharomyces* and *Hanseniaspora* during wine fermentation (Langenberg et al., 2017), may similarly facilitate horizontal transfer.

The mitochondrial genomes generated in this study provide many opportunities to further our understanding of evolution beyond the scope of this study. Pairing the data with the previously generated nuclear genomes will help elucidate the interplay between these two genomes. Interactions between mitochondrial and nuclear loci (mito-nuclear epistasis) have been demonstrated to affect phenotypic variation in yeasts (Paliwal et al., 2014; Nguyen et al., 2020b, 2023; Visinoni and Delneri, 2022; Biot-Pelletier et al., 2023) and a diverse array of model systems (Dowling et al., 2007; Burton and Barreto, 2012; Mossman et al., 2016). For many existing mitochondrial genomes, any analysis of such interactions was previously often complicated by the lack of a corresponding nuclear genome or by mismatches between the strains sequenced for a given species. By mining most of this new mtDNA dataset from a dataset of high-quality nuclear genomes (Shen et al., 2018), many of these previous limitations have been lifted, which has already enabled the novel insights described here. The breadth and the richness of these paired nuclear-mitochondrial datasets promise to greatly accelerate research into the evolution of yeast mitochondrial genomes.

## 4 Materials and Methods

### 4.1 Mitochondrial Genome Rescue, Assembly, and Annotation

We searched 332 yeast genome assemblies for mitochondrial contigs using a two-pronged, reference-based approach (Shen et al., 2018). First, we curated a set of reference mtDNAs from all accessions in Genbank matching Saccharomycotina and with the source as “mitochondrion” in September 2018 to generate a set of 110 published mtDNAs with a single representative per species (Supplemental Table 1). Existing annotations were curated based on length and presence of stop codons, and they were renamed for consistent formatting. When annotations were not available, new annotations were generated using MFANNOT (Lang et al., 2007). We identified putative mitochondrial contigs based on two BLAST strategies searches (v2.8.1). First, the coding sequences (CDS) from the curated references were used as queries to search each assembly, and contigs with at least 10 hits >70% coverage and e-value <0.001 were retained. Second, the contigs from each assembly were used as queries against the complete reference mtDNAs, and contigs with at least one 25% coverage hit with e-value <0.001 were retained. These contigs were then preliminarily annotated using MFANNOT (Lang et al., 2007) to estimate gene content (Lang et al., 2007). To eliminate contigs that were likely short duplicates of mitochondrial sequences transferred to the nuclear genome, also known as NUMTs (Hazkani-Covo et al., 2010; Xue et al., 2023), we filtered out contigs that did not possess at least one mitochondrial gene per 20 kb. Contigs larger than 300kb were also removed to eliminate any complete mtDNA duplicates in large nuclear contigs.

Assembly methods for nuclear genomes are generally not optimized for mitochondrial sequences, so we reassembled genomes for which sequencing reads were readily available, including 196 species sequenced in Shen et al. 2018 and 92 additional species included in that dataset that we resequenced to replace an older nuclear assembly as part of the Y1000+ Project (Supplemental Table 1) (Opulente et al., 2023). Reassembly was done using either plasmidSPAdes v3.9.0 (Antipov et al., 2016) or NOVOPlasty v4.2 (Dierckxsens et al., 2017). We annotated these assemblies using MFANNOT (Lang et al., 2007) and then searched for mitochondrial contigs as described above. For NOVOPlasty, multiple assemblies were constructed using different seeds either using the genes found in the putative mitochondrial contigs extracted from the nuclear assembly or the CDS from the closest available genome based on the nuclear phylogeny in the curated reference set. The putative mitochondrial contigs isolated from the nuclear assembly and the mitochondrial reassemblies were assessed based on completeness (% of expected genes present, excluding *RPS3* and Complex I genes when none were present), contiguity (% of genes found on each contig), and circularity (count of reads that map across the contig endpoints after shifting the sequence such that the original breakpoint is internal in the permuted contig), and a single assembly was chosen for each species. We prioritized completeness and used contiguity and circularity to break ties. Generally, NOVOPlasty performed best, followed by plasmidSPAdes, while the existing contigs from the nuclear assembly were best in a small minority of cases. For the final dataset, we combined these assemblies with the curated reference set, retaining one assembly per species and choosing the reference assembly for a species when available.

All new assemblies, as well as existing mtDNAs that were not annotated, were annotated from scratch; all genome annotations, including published ones, were curated for consistency and to improve accuracy as described below. The translation table for each species was estimated using codetta v2.0 (Shulgina and Eddy, 2021). Yeast mitochondrial translation tables fall into either the Mold, Protozoan, and Coelenterate Mitochondrial Code and the Mycoplasma/Spiroplasma Code (NCBI table 4, hereafter referred to as the fungal code), which is consistent with other fungi, or the yeast mitochondrial code (NCBI table 3, originally based on *S. cerevisiae,* hereafter referred to as the *Saccharomyces* code) based on additional reassignments of AUA and CUN codons, which typically define the order Saccharomycetales. The exact placement of this transition was difficult to determine due to a loss of CUN codons in many Saccharomycetales, particularly *Kluyveromyces* species and other closely related genera. In many species, the CUN reassignment is supported by codetta, but the AUA reassignment is not, and the modified tRNA required for this reassignment is not present, which is consistent with a previous analysis of codon usage among Saccharomycotina mtDNAs (Christinaki et al., 2022). Currently, no translation table exists for the CUN reassignment without the AUA reassignments, so we used the *Saccharomyces* code when the CUN reassignment was supported and the fungal code for all others (Table S1). The AUA reassignment in the *Saccharomyces* code allows for this codon to be treated as a start codon by MFANNOT, which resulted in many misannotations at the 5’ end of genes. No examples of AUA being used as a valid start codon in yeasts have been described. We rectified this issue by reannotating all assemblies using table 4 to define start and end coordinates; we then used the *Saccharomyces* code for translation when appropriate. Finally, all annotations (for new and existing mtDNAs) were further manually curated to eliminate truncated genes, annotations split across contigs, and annotations containing large extensions due to misannotated introns or readthroughs. We identified several *Kazachstania* species with frameshifts consistent with the +1C frameshift mechanism previously described (Szabóová et al., 2018). To match the formatting in GenBank for those references, we encoded these as single-bp introns, but these were excluded from all intron analyses. We did not observe any *byp* elements, as described in *Magnusiomyces tetraspermus,* in the coding sequences of other species (Lang et al., 2014).

### 4.2 Mitochondrial Phylogeny Construction

We determined phylogenetic relationships among mitochondrial genomes based on the core set of genes shared by all species: *COX1*, *COX2*, *COX3*, *COB*, *ATP6*, *ATP8*, and *ATP9*. Complex I genes were excluded due their loss in a large fraction of the species. Protein sequences were aligned for each gene using MAFFT using the E-INS-I option (Katoh and Standley, 2013), and CDS were

codon-aligned using the protein alignment. The alignments were concatenated and then filtered to retain only sites in which 95% of sequences were not gaps using trimAl (Capella-Gutiérrez et al., 2009). We built multiple phylogenies from the filtered alignment using IQ-TREE using the mitochondrial substitution model (Minh et al., 2020). These phylogenies were highly concordant, except for the placement of the fast-evolving *Hanseniaspora* lineage. The topology most consistent with the nuclear phylogeny was selected as the final tree. Phylogenetic correction of correlations of genome size versus GC content were done using a generalized least squares approach (using gls from nlme (Pinheiro J, Bates D, 2023)) using a co-variation matrix generated using a Brownian motion model (using corPagel from ape (Paradis and Schliep, 2019)).

### 4.3 Estimating Patterns of Selection

To investigate patterns of selection on mitochondrial genes, we split the phylogeny into smaller groups at roughly the genus level to avoid saturation of synonymous substitutions (Table 1). For each of the genes in the core set, we built subtrees for each group and estimated ω along each branch of the subtree using PAML under model 1 (allowing variable ω for each branch) (Yang, 2007). For each gene, the ω value was determined as the mean of the values for all branches in the subtree for which there were sufficient synonymous substitutions (dS > 0.01).

### 4.4 Evaluating Evidence for Horizontal Transfer of Mitochondrial Introns

Possible HGTs of mitochondrial introns were determined based on an all-versus-all BLAST of mitochondrial introns against each other. Mitochondrial introns among closely related species are expected to share limited sequence similarity due to poor conservation of non-coding sequences, though elements that contribute to intron splicing may be under purifying selection. Thus, we set a conservative threshold that the bit score of each hit must be at least 50% of the maximum possible bit score determined by the self-to-self comparison of each intron and have an e-value < 10^-10^. Shared relationships within groups are likely to be due to vertical descent, although there is evidence that HGT frequently occurs at this scale (Wu and Hao, 2014), but such high sequence similarity at large phylogenetic distances is likely due to HGT. Clustering of intron sequences was performed using the Louvain method (Blondel et al., 2008) implemented in the igraph (Csárdi et al., 2023) package of R.

## 6 Author Contributions

All authors assisted in preparation of the final manuscript. JFW designed and implemented research, performed all computational and statistical analyses, managed data, and prepared the figures. ALL assisted in developing the methodology for isolating mitochondrial contigs from existing whole genome assemblies. DAO led genome sequencing for all resequenced genomes from Shen et al. 2018. AR and CTH designed the research, obtained funding, and supervised the project.

## 8 Acknowledgments

We thank Jacob L. Steenwyk, Xiaofan Zhou, Trey K. Sato, Hittinger Lab members, and Y1000+ Project members for helpful comments; the University of Wisconsin Biotechnology Center DNA Sequencing Facility (Research Resource Identifier – RRID:SCR_017759) for providing DNA sequencing facilities and services; Wisconsin Energy Institute staff for computational support; and the Center for High-Throughput Computing at the University of Wisconsin-Madison (https://doi.org/10.21231/GNT1-HW21).

## 9 Supplementary Material

Supplementary Material should be uploaded separately on submission, if there are Supplementary Figures, please include the caption in the same file as the figure. Supplementary Material templates can be found in the Frontiers Word Templates file.

Please see the Supplementary Material section of the Author guidelines for details on the different file types accepted.

## 12 Data Availability Statement

All supporting data and analyses are available at https://figshare.com/s/9266509ee3a167725b5f. This link will be replaced with a public link on acceptance.

Supplemental Figure 1. Mitochondrial Genome Quality.

The completeness (proportion of expected genes present on the best contig, excluding *RPS3*, and excluding *NAD* genes when none were present in the assembly) and contiguity (proportion of genes found present on the best contig) of putative mitochondrial contigs are shown for A) public genome assemblies included in (Shen et al., 2018), B) newly sequenced genome assemblies included in (Shen et al., 2018), and C) our final mitochondrial genome dataset. Genomes with high completeness but lower contiguity were typically well represented by only two contigs.

Supplemental Figure 2. Genome size versus GC Content.

The correlation between genome size and GC content is indicated with individual genomes labeled by taxonomic order as in Figure 3. Larger genomes tended to have lower GC content, but the correlation was only weakly significant. Phylogenetic correction increased the strength of the correlation, but it was no longer statistically significant. GC content does not appear to play a central role in influencing genome size.

Supplemental Figure 3. Candidates for Intron HGT across Taxonomic Orders.

Intron sequences were compared using BLAST, and scores were used to generate clusters of closely related introns. Four clusters showed high homology between introns from different taxonomic orders; three are displayed here, while the fourth one is in Figure 6C. Introns are labeled by taxonomic order as in Figure 3.

