## Supplementary material for "Mitochondrial Genome Diversity across the Subphylum Saccharomycotina": Figures S1-S3

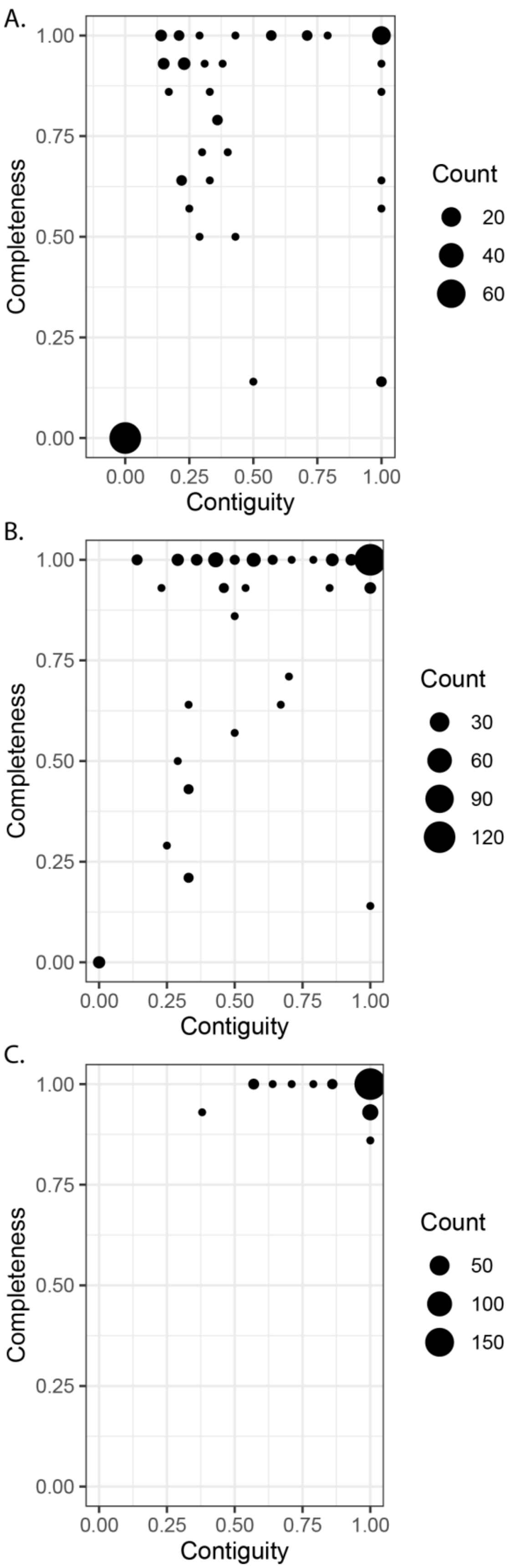


Supplemental Figure 1 Mitochondrial Genome Quality.
The completeness (proportion of expected genes present on the best contig, excluding *RPS3*, and excluding *NAD* genes when none were present in the assembly) and contiguity (proportion of genes found present on the best contig) of putative mitochondrial contigs are shown for A) public genome assemblies included in (Shen et al., 2018), B) newly sequenced genome assemblies included in (Shen et al., 2018), and C) our final mitochondrial genome dataset. Genomes with high completeness but lower contiguity were typically well represented by only two contigs.

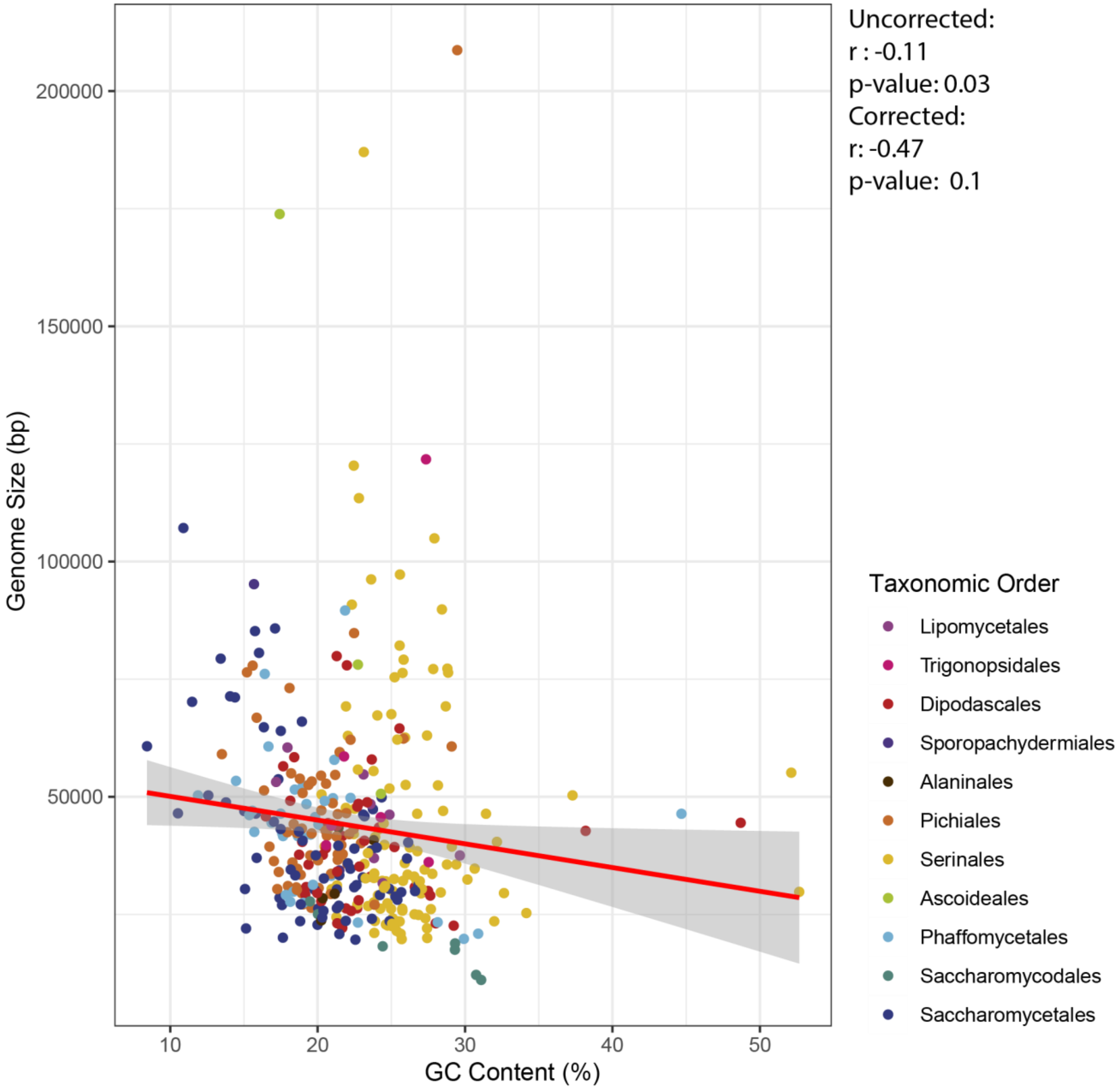


Supplemental Figure 2 Genome size versus GC Content.

The correlation between genome size and GC content is indicated with individual genomes labeled by taxonomic order as in Figure 3. Larger genomes tended to have lower GC content, but the correlation was only weakly significant. Phylogenetic correction increased the strength of the correlation, but it was no longer statistically significant. GC content does not appear to play a central role in influencing genome size.

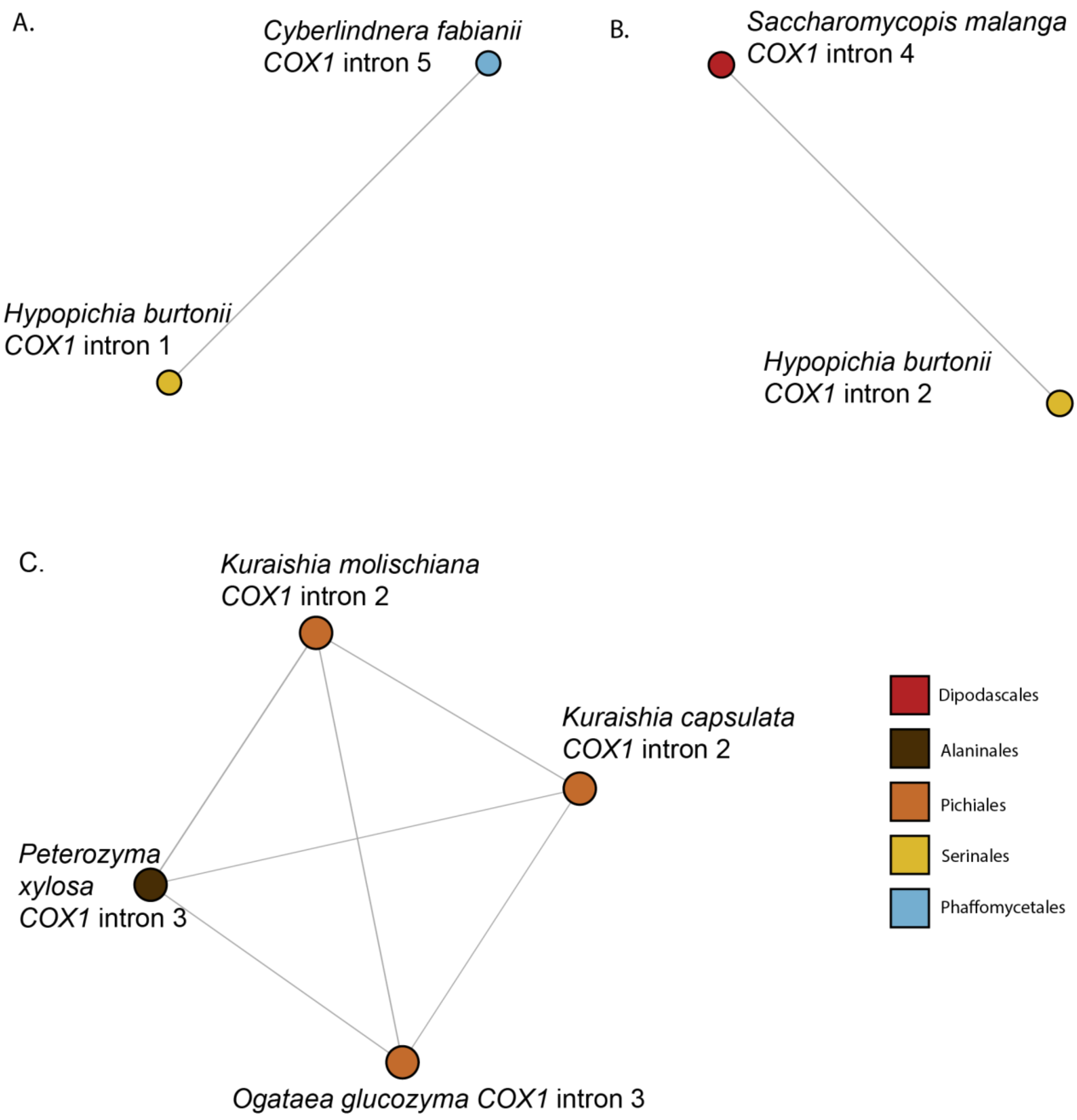


Supplemental Figure 3. Candidates for Intron HGT across Taxonomic Orders.

Intron sequences were compared using BLAST, and scores were used to generate clusters of closely related introns. Four clusters showed high homology between introns from different taxonomic orders; three are displayed here, while the fourth one is in Figure 6C. Introns are labeled by taxonomic order as in Figure 3.
